# *KCNH1* variants associated with neurodevelopmental disorders exhibit prominent gain-of-function

**DOI:** 10.64898/2026.02.09.704863

**Authors:** Christopher H. Thompson, Christine Q. Simmons, Isabella K. He, Nirvani Jairam, Reshma R. Desai, Esther Cha, Alexandra E. Hong, Jean-Marc DeKeyser, Linda C. Laux, Michaelle Jinnette, Carlos G. Vanoye, Alfred L. George

**Affiliations:** Department of Pharmacology, Feinberg School of Medicine, Northwestern University, Chicago, IL 60611 USA; Ann and Robert H. Lurie Children’s Hospital, Chicago, IL 60611 USA; Cure KCNH1 Foundation, Encinitas, CA 92024 USA

## Abstract

Pathogenic variants in *KCNH1*, encoding the human ether-á-go-go voltage-gated potassium channel (EAG1, K_V_10.1), are associated with the neurodevelopmental disorders Temple-Baraitser syndrome (TBS), Zimmermann-Laband syndrome (ZLS), and epileptic encephalopathy. Investigating the functional consequences of *KCNH1* variants can provide information about molecular and cellular pathophysiological mechanisms, and help guide therapeutic directions. In this study, we determined the functional properties of 26 disease-associated *KCNH1* variants including three recurrent and five that were previously unpublished. Findings were compared to the wildtype (WT) channel along with six rare population variants. Our approach used automated patch clamp recording of heterologously expressed *KCNH1* variants in both the homozygous state (variant alone) and the clinically-relevant heterozygous state (variant co-expressed with WT channels). We also examined the impact of alternative splice isoforms on functional consequences of specific variants. We demonstrated that the majority of disease-associated *KCNH1* variants exhibit features consistent with gain-of-function including hyperpolarized voltage-dependence of activation, accelerated activation kinetics, or both that promote hyperpolarization of the resting membrane potential. By contrast, one unique variant discovered previously in an autism cohort had a depolarized voltage-dependence of activation and was associated with a depolarized resting potential in cells. Pharmacological testing of *KCNH1* variants demonstrated differences in potency for channel block by imipramine and an investigational non-selective potassium channel blocker (LY97241) for specific variants. Collectively, our findings support gain-of-function as the most common, albeit not exclusive, molecular mechanism underlying *KCNH1*-related disorders and suggest opportunities for potential therapeutic drug repurposing strategies for these conditions.

## Introduction

Channelopathies of the central nervous system are frequently discovered genetic etiologies of neurological and neurodevelopmental disorders.^1^ Pathogenic variants in several genes encoding voltage-gated ion channels are often reported in case series of early-onset epilepsy and related neurodevelopmental disorders.^2-6^ Investigating the molecular basis for these disorders has provided a rich source of new knowledge informing about disease mechanisms and guiding therapeutic strategies.^7-10^

One recently discovered channelopathy gene (*KCNH1*) associated with complex and severe neurodevelopmental disorders encodes the human ether-á-go-go (EAG1, K_V_10.1) voltage-gated potassium channel widely expressed in brain including both excitatory and inhibitory neurons.^11^ Heterozygous *KCNH1* variants were first described in association with Temple-Baraitser syndrome (TBS) and Zimmermann-Laband syndrome (ZLS),^12,13^ which exhibit overlapping clinical features including hypotonia, developmental delay, intellectual disability, seizures, and variable presence of gingival enlargement, facial and digit dysmorphism, hypoplastic or absent thumb and toe nails. Subsequently, there were independent reports of several novel and recurrent *KCNH1* variants associated with TBS and ZLS as well as in cases without all manifestations of these disorders.^14-24^ Most reported *KCNH1* variants are *de novo* and associated with poor long-term neurodevelopmental outcomes.

Efforts to determine the functional consequences of *KCNH1* variants have been conducted *in vitro* using a variety of heterologous expression systems including *Xenopus* oocytes, and cultured mammalian cells (HEK293, Chinese hamster ovary [CHO] cells). The functional properties of seven missense variants and one compound genotype with two missense variants occurring in *cis* have been reported.^12,13^ When compared to the wildtype (WT) channel, most variants exhibit a hyperpolarized voltage-dependence of activation and variable effects on channel gating kinetics that have been interpreted as gain-of-function. However, these studies mainly examined channel variants expressed in the homozygous state (e.g., variant expressed alone) rather than the more clinically relevant heterozygous state (e.g., variant co-expressed with WT). Surprisingly, the functional effects of the most recurrent *KCNH1* variant (R357Q) have not been reported nor have there been published studies investigating rare missense variants discovered in the general population. These gaps in knowledge limit our understanding of the full pathophysiological spectrum of *KCNH1*-related disorders.

In this study, we conducted a systematic investigation to determine the functional consequences of approximately half of all reported disease-associated *KCNH1* variants including five previously unpublished variants and six rare missense population variants. We employed high throughput automated patch clamp recording to enable more robust sampling without operator bias in cell selection, examined variants in both the homozygous and heterozygous states, and investigated the impact of alternative *KCNH1* mRNA splicing on variant functional effects. We determined that most variants exhibit gain-of-function properties in the heterozygous state and drive the resting membrane potential of cells to more negative voltages. Our work contributes to emerging ideas about the pathophysiology of *KCNH1*-related disorders and may guide development of new therapies.

## Results

### KCNH1 variant landscape and genotype-phenotype correlations

We ascertained the *KCNH1* variant landscape from multiple sources. We collated literature-reported *KCNH1* variants annotated in the Human Gene Mutation Database (HGMD),^30^ and distinct variants registered in ClinVar.^31^ An additional cohort of deidentified participants in the KCNH1 International Registry established by the Cure KCNH1 Foundation was examined. Additional population variants with minor allele frequencies greater than 0.00005 were identified in the Genome Aggregation Database (gnomAD).^32^ Variant codons were numbered based on the amino acid sequence predicted from KCNH1 transcript isoform 1 (NCBI RefSeq NM_172362). Approximately 50 known disease-associated variants have been reported in the literature with a few notable recurrent variants discovered in unrelated probands.

We selected 6 nonsynonymous population variants and 26 disease-associated missense *KCNH1* variants for functional testing. **Table 1** presents a summary of clinical features associated with *KCNH1* variants investigated in this study and **Fig. 1** illustrates the distribution of variants in the KCNH1 protein. We included three recurrent variants (R357Q, L489F, G496R), and five previously unpublished variants (N238D, G375R [c.1123G>C], A492S, R520W, P560S). Most variants were associated with overlapping clinical features consistent with TBS or ZLS including hypotonia, developmental delay, intellectual disability, seizures, and variable presence of gingival enlargement, facial and digit dysmorphism, abnormal thumb and great toe nails (hyponychia or anonychia). *KCNH1* variants associated with less typical syndromes included R520W associated with a prominent seizure phenotype and normal muscle tone, V569M identified in an autism cohort,^17^ and V713E discovered in two probands with epilepsy but without other features of TBS or ZLS.^19^ One individual heterozygous for V713E inherited the variant from an unaffected parent, and in the other individual this variant was somatic in brain tissue resected for treatment of refractory epilepsy. In addition, V713E is classified as likely benign in ClinVar, whereas most other disease-associated variants we studied were classified as pathogenic or likely pathogenic (**Table 1**). Two disease-associated variants (R520Q, V569M) occurred infrequently in the gnomAD database, whereas all others were absent in gnomAD.

**Table 1.**
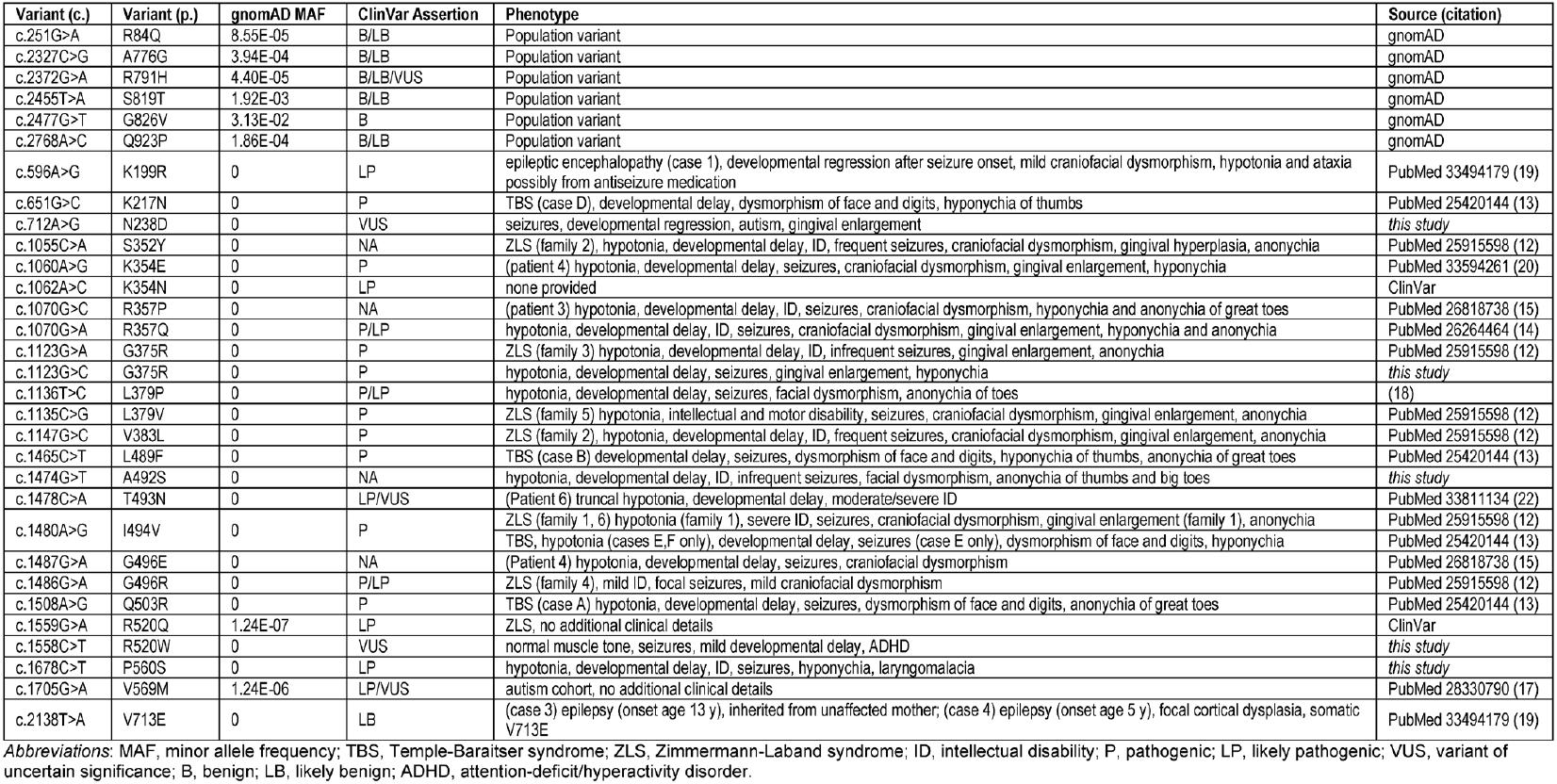
Clinical and genetic features associated with *KCNH1* variants.

**Fig. 1.**
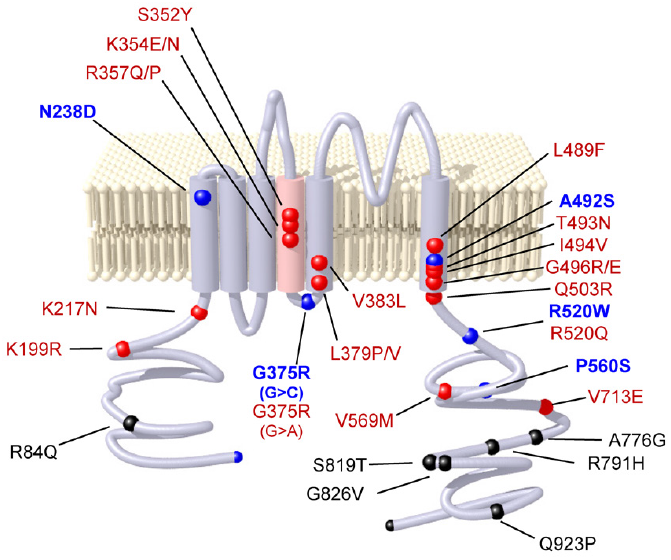
Location of KCNH1 variants studied. The locations of all studied variants are illustrated in a simplified protein structure. Black spheres represent population variants, red spheres represent previously reported disease-associated variants, and blue sphere represent previously unpublished disease-associated variants. For variants affecting the same codon, both amino acid substitutions are indicated (e.g., K354E/N) but only one symbol color is visible.

### Functional consequences of KCNH1 variants

We investigated the functional consequences of KCNH1 variants expressed in HEK293T cells using automated patch clamp recording. Initial experiments were conducted using cells expressing only WT or variant KCNH1 (homozygous state), and subsequently each variant was co-expressed with WT KCNH1 to mimic the heterozygous state observed in affected individuals. We analyzed data from a total of 6,821 recorded cells. For heterozygous experiments, we quantified the level of successful co-transfection (86.1 ± 1.1% [95%CI]) using flow cytometry to detect distinct fluorescent proteins coupled to expression of WT or variant plasmids. Cells expressing WT KCNH1 exhibited a slowly activating, non-inactivating outward current (**Fig. 2A**). In the homozygous state, all variants except G496E, G496R and R520W exhibited levels of current density larger than endogenous currents although there was substantial variant-to-variant variability (**Supplementary Fig. S1**). For clinical relevance, most further analyses were done with variants expressed in the heterozygous state.

**Figure 2.**
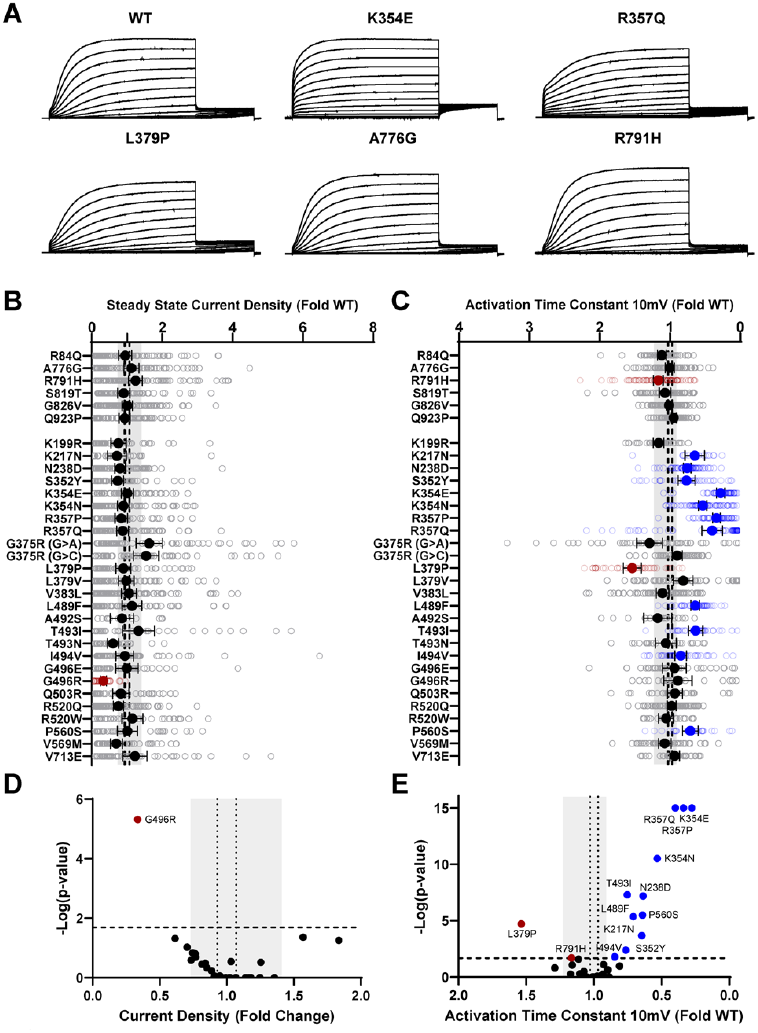
Disease-associated KCNH1 variants exhibit altered activation kinetics. (**A**) Average whole-cell current traces from cells co-expressing WT, three disease-associated variants (K354E, R357Q, L379P), and two population (A776G, R791H) KCNH1 variants in the heterozygous state. (**B**) Average deviation of steady-state current density from cells co-expressing WT and either population or disease-associated KCNH1 variants. (**C**) Average deviation from WT of the activation time constant measured at 10 mV for population and disease-associated variants. (**D**,**E**) Volcano plots highlighting variants significantly different from WT for steady-state current density in panel **D** and activation time constant in panel **E**. In volcano plots, vertical dashed lines denote the 95% CI for WT, while the vertical gray bar denotes the range of 95% CI for the population variants. Blue symbols denote variants with significantly faster activation kinetics (time constants < 1.0) and red symbols denote variants with significantly slower activation (time constants > 1.0) at a statistical threshold of *P* < 0.02 (*n* = 15-94).

We compared steady state current density elicited with a test depolarization to +50 mV and compared heterozygous variants to WT channels (**Supplementary Fig. 2**). None of the population variants had current densities significantly different from WT (**Fig. 2B,D**). Among disease-associated variants, only G496R had a significant difference (65% smaller) in current density compared to WT (**Fig. 2B,D**).

Activation kinetics quantified as a single time constant were determined at a test potential of +10 mV. Most population variants exhibited WT-like activation kinetics (**Fig. 2**) with the exception of R791H, which activated more slowly (e.g., larger time constant) than WT (**Fig. 2A,C,E**). Eleven of the 26 disease-associated variants (K217N, N238D, S352Y, K354E, K354N, R357P, R357Q, L489F, T493I, I494V, P560S) exhibited faster activation compared to WT (**Fig. 2C,E**). For some disease-associated variants, activation onset was rapid as exemplified by K354E or had initial instantaneous activation as illustrated for R357Q (**Fig. 2A**; **Supplementary Fig. S2**). Only one disease-associated variant (L379P) had slower activation kinetics compared to WT channels, which was outside the range of variation observed for the population variants (**Fig. 2C,E**). A different mutation of the same codon (L379V) exhibited no difference in activation kinetics.

We compared the voltage-dependence of activation for population and disease-associated variants in the heterozygous state to WT channels. While none of the population variants showed any divergence from WT (**Fig. 3A,B**), 17 of the 26 disease-associated variants exhibited a hyperpolarized voltage-dependence of activation compared to WT (**Fig. 3A-C**). The most recurrent variant, R357Q, showed a modest degree (-4.4 mV) of activation V_1/2_ hyperpolarization compared to WT channels (denoted as ΔV_1/2_), while other variants exhibited much larger degrees of hyperpolarization (e.g., K354E, ΔV_1/2_: -12.9 mV). Only V569M had a depolarized (ΔV_1/2_: +10.8 mV) voltage-dependence of activation relative to WT.

**Figure 3.**
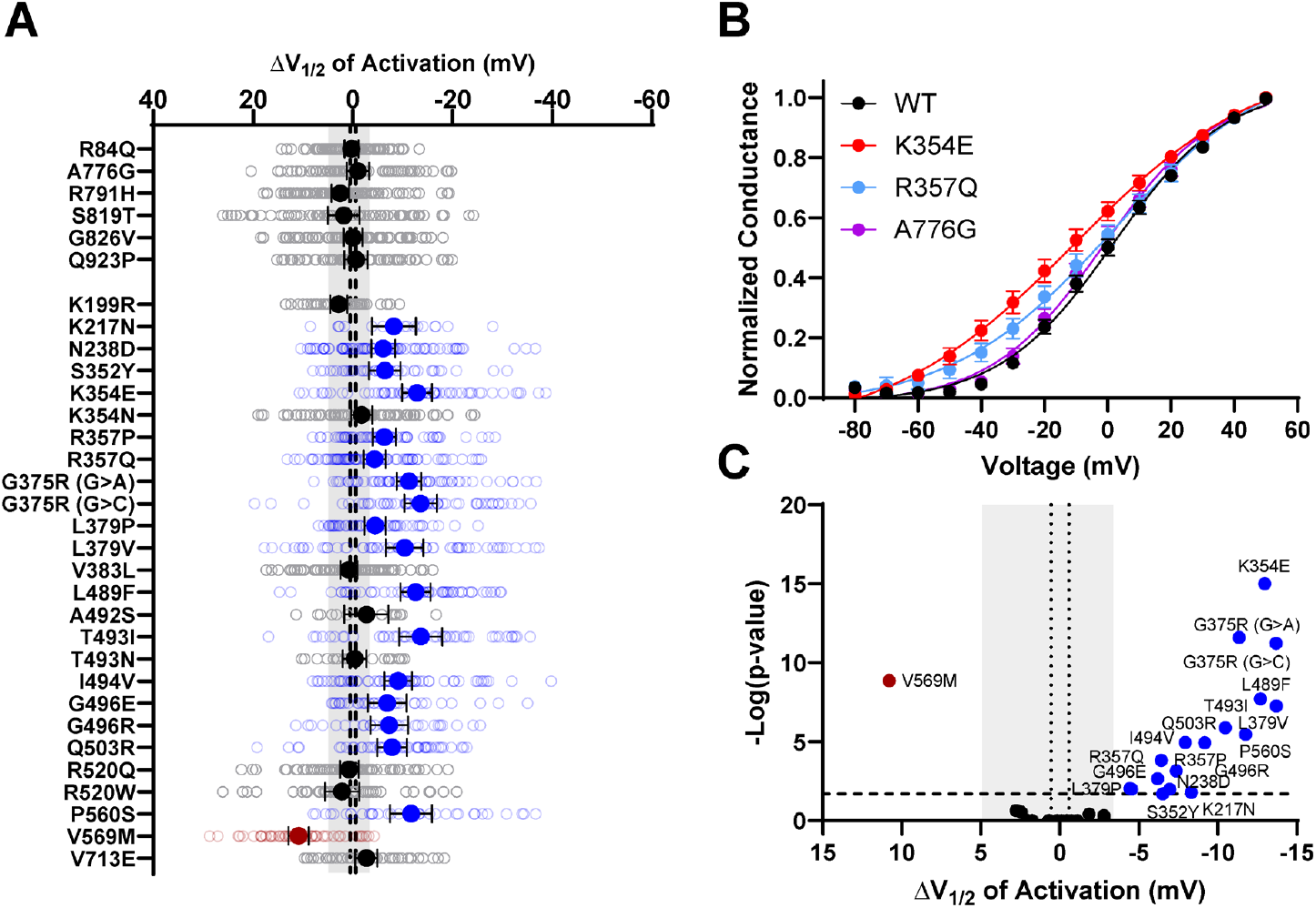
Disease-associated KCNH1 variants have altered voltage-dependence of activation. (**A**) Average deviation from WT in voltage-dependence of activation for population or disease-associated variants in the heterozygous state. (**B**) Average conductance-voltage relationships for WT, two disease associated variants, K354E, R357Q, and one population variant, A776G. (**C**) Volcano plots highlighting variants significantly different from WT for voltage-dependence of activation. Vertical dashed lines denote the 95% CI for WT, while the vertical gray bar denotes the range of 95% CI for the population variants. Blue symbols denote variants with significantly hyperpolarized activation and red symbols denote variants with significantly depolarized activation at a statistical threshold of *P* < 0.02 (*n* = 15-94).

Not all variants exhibited significant dysfunction. Among the 26 disease-associated KCNH1 variants studied, two (K199R, V713E) showed no significant differences for any measured parameters compared to WT channels when expressed in either the homozygous or heterozygous state. Three other variants (V383L, A492S, T493N) exhibited significant effects only in the slope factor determined from voltage-dependence of activation curves when expressed in the homozygous state, but no significant difference from WT in the heterozygous state. Lastly, two variants affecting the same codon (R520Q, R520W) exhibited significant effects on multiple parameters in the homozygous state but none in the heterozygous state.

### Expression of KCNH1 splice isoforms in human cortex

Because some disease-associated variants exhibited limited or no significant functional differences, we considered whether alternative forms of KCNH1 may be more relevant. KCNH1 mRNA transcripts undergo alternative splicing resulting in two major splice isoforms, a long isoform (isoform 1) consisting of 11 exons encoding a 989 amino acid protein, and a shorter isoform (isoform 2) that excludes 81 bp of exon six resulting in a 962 amino acid protein having a shorter S3-S4 loop.^33^ We used next generation sequencing to quantify the contribution of each splice isoform to total KCNH1 mRNA expression in human brain and investigated whether this alternative splicing event is developmentally regulated. We examined KCNH1 splice isoform expression in human cerebral cortex obtained from brain bank donors ranging in age from 18 post-conception weeks to 20 years. Across all age groups tested, both isoforms are expressed with isoform 2 being approximately twice as abundant. (**Fig. 4A**; **Supplementary Fig. S3**). Isoform 1 had a slightly larger contribution to KCNH1 total transcript during childhood compared to the prenatal period (**Fig. 4B**). However, noting that both isoforms are expressed in abundance at all age groups, KCNH1 alternative splicing does not appear to be regulated developmentally.

**Figure 4.**
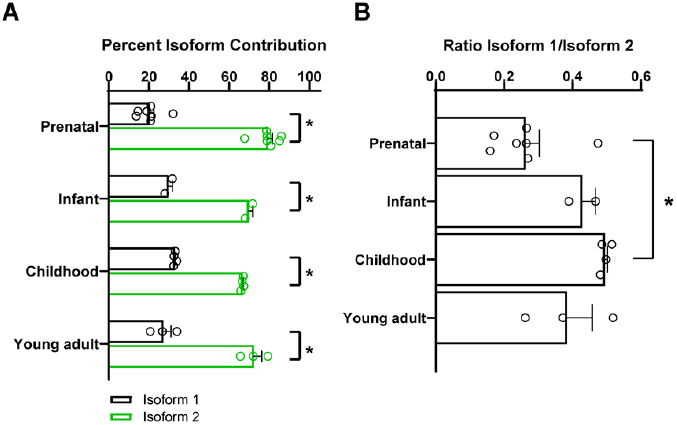
Expression of KCNH1 alternate splice isoforms in human brain. (**A**) Percent contribution of each splice isoform to total KCNH1 mRNA from human brain samples. (**B**) KCNH1 isoform 1/isoform 2 ratio across age groups. Each age group is represented by 2-7 samples. Data are mean ± SEM. The asterisk represents *P* < 0.05.

### Effect of KCNH1 alternative splicing on disease associated variants

We compared the functional properties of WT KCNH1 in the context of the two splice isoforms. KCNH1 isoform 2 produces slightly larger whole-cell currents with a more depolarized voltage-dependence of activation and slower activation kinetics compared to isoform 1 (**Supplementary Fig. S4A-D**). We compared the functional properties of the most recurrent variant (R357Q) in the two splice isoforms and observed qualitatively similar but quantitatively different effects in the homozygous state (**Supplementary Fig. S4E**). In particular, the voltage-dependence of activation observed for R357Q was hyperpolarized relative to WT in both isoforms, but the difference was larger in isoform 2 (isoform 1: ΔV_½_ = -13.7 mV; isoform 2: ΔV_½_ = -24.6 mV). Similarly, cells expressing A492S in isoform 2 exhibited a significant difference in activation V_½_ in both homozygous (ΔV_½_ = -29.5 ± 4.9 mV, *P*<0.0001) and heterozygous (ΔV_½_ = -17.5 ± 2.9 mV, *P* <0.0001) states, whereas the differences observed in isoform 1 were not statistically significant (homozygous: ΔV_½_ = -6.0 ± 6.4 mV, *P*>0.99; heterozygous: ΔV_½_ = -2.8 ± 4.0 mV, *P* =0.82) (**Fig. 5**). By contrast, two variants without any functional differences in isoform 1 (K199R, V713E) exhibited no differences in any functional parameter in isoform 2 in the homozygous state and therefore were not tested in the heterozygous state. The three variants (V383L, A492S, T493N) with limited functional effects in isoform 1 in the heterozygous state, exhibited significant differences relative to WT channels when expressed in isoform 2 in the heterozygous state (**Fig. 5**; **Supplementary Fig. S5**). Lastly, V569M, which is the only variant with a depolarized voltage-dependence of activation V_1/2_, exhibited similar effects in isoform 2 in the heterozygous state and had a significantly smaller peak current density (**Fig. 5**; **Supplementary Fig. S5**). These findings indicate that some *KCNH1* variants exhibit isoform-dependent dysfunction.

**Figure 5.**
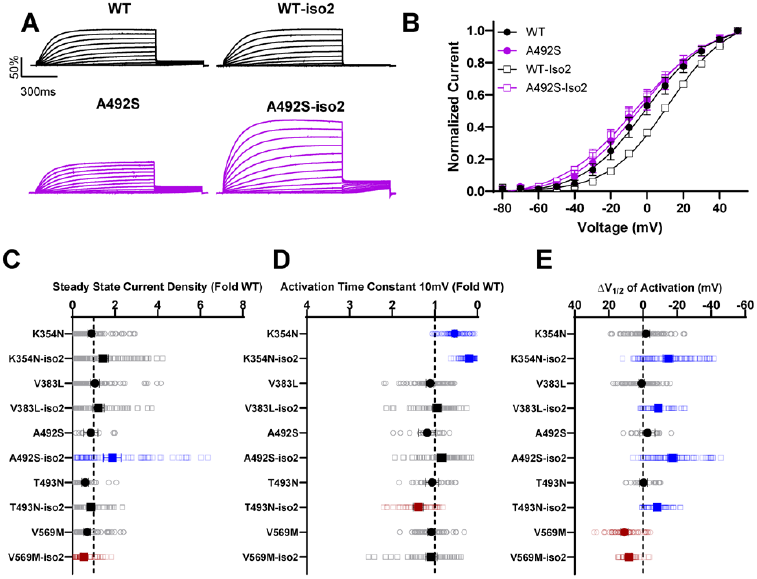
Splice isoform impacts functional properties of KCNH1 variants. (**A**) Average whole-cell current traces from cells co-expressing WT and A492S in both KCNH1 splice isoforms in the homozygous state. (**B**) Average conductance-voltage relationship of WT and A492S in both splice isoforms. (**C-E**) Average deviation from WT in the heterozygous state for steady-state current density in panel **C**, activation time constant relative to WT in panel **D**, and difference in voltage-dependence of activation (ΔV_1/2_) in panel **E**. Blue and red symbols denote variants significantly different from WT at a statistical threshold of *P* < 0.02 (*n*=14-107).

### Impact of KCNH1 variants on resting potential

KCNH1 is a subthreshold potassium channel that helps stabilize the resting membrane potential (RMP) of neurons.^34,35^ In current clamp experiments, we observed that expression of WT KCNH1 alone hyperpolarized the RMP of HEK293T cells to -61.4 ± 0.8 mV compared to - 17.5 ± 3.1 mV recorded from non-transfected cells. Resting membrane potential recording from cells expressing population variants was not significantly different from WT-expressing cells. We hypothesized that a hyperpolarized voltage-dependence of activation or faster activation will enable larger potassium conductance at negative membrane potentials with greater impact on the RMP. Consistent with our hypothesis, heterozygous expression of 21 of the 26 disease-associated variants induced significantly greater RMP hyperpolarization compared to WT (**Fig. 6A,B**). A492S and T493N only showed hyperpolarization of RMP when expressed in isoform 2, while V569M depolarized RMP only when expressed in isoform 2 (**Fig. 6B**). However, some variants with abnormal functional properties only in the homozygous state (e.g., R520Q, R520W) did not affect the RMP in the heterozygous state.

**Figure 6.**
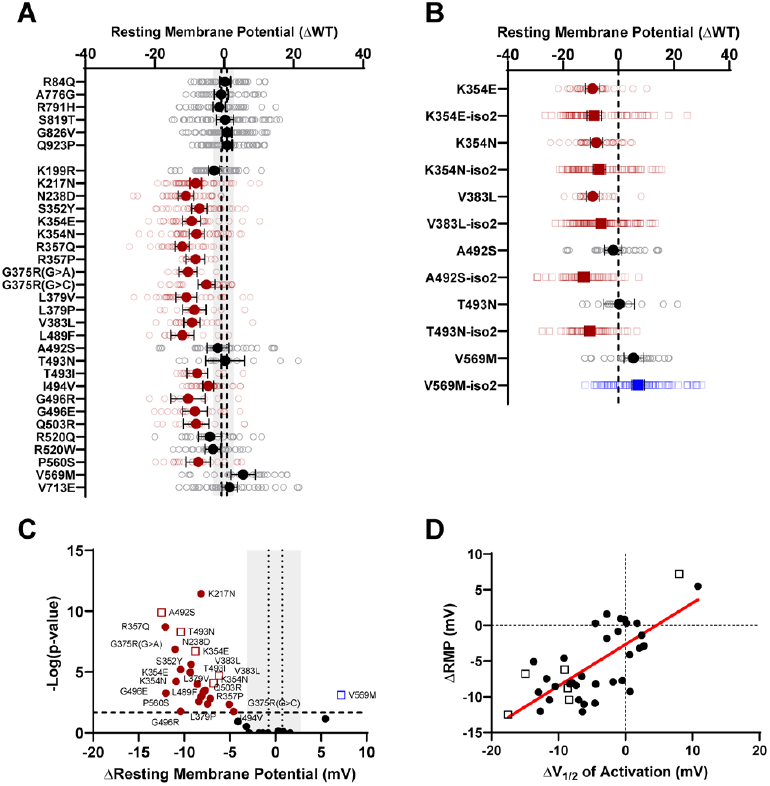
Disease-associated KCNH1 variants hyperpolarize resting membrane potential. (**A**) Average difference in resting membrane potential (ΔRMP) between cells expressing WT KCNH1 compared to cells co-expressing WT and variants studied in isoform 1. (**B**) Average difference in resting membrane potential (ΔRMP) between cells expressing WT compared to cells co-expressing WT and selected variants in KCNH1 isoform 1 (circles) or isoform 2 (squares). (**C**) Volcano plots highlighting variants significantly different from WT for resting membrane potential. Vertical dashed lines denote the 95% CI for WT, while the vertical gray bar denotes the range of 95% CI for the population variants. (**D**) Correlation between ΔRMP and ΔV_1/2_ of activation. Red line is a fitted regression line (*P*<0.0001). Red symbols denote significantly hyperpolarized RMP and blue symbols denote significantly depolarized RMP at a statistical threshold of *P* < 0.02 (*n*=9-67).

The difference in voltage-dependence of activation from WT was well correlated with the degree of RMP hyperpolarization (**Fig. 6D**). By contrast, activation kinetics were only modestly correlated with the degree of RMP hyperpolarization (**Supplementary Fig. S6**). We concluded that the shift in activation voltage-dependence was the primary driver of RMP hyperpolarization. This is consistent with our observation that KCNH1 variant G496R was also associated with RMP hyperpolarization even though current in the heterozygous state was low.

### Drug repurposing for KCNH1 gain-of-function variants

Because the majority of disease-associated KCNH1 variants cause gain-of-function, we speculated that there may be existing drugs or investigational compounds that can suppress channel hyperactivity and offer potential therapeutic benefit. Several approved drugs with good brain penetration used to treat psychiatric conditions including chlorpromazine, haloperidol and imipramine have been reported to inhibit KCNH1 channels *in vitro*.^36-39^ Additional drugs such as astemizole, and mibefradil are also potent KCNH1 inhibitors,^36,40,41^ but were withdrawn from the market due to proarrhythmic effects mediated by block of the human ether-á-go-go related gene (hERG encoded by *KCNH2*) in heart. An investigational compound (LY97241) and the antiarrhythmic drug clofilium from which it was derived are non-selective K+ channel blockers with high potency for KCNH1.^42^

As a small-scale proof-of-concept drug repurposing study, we quantified the KCNH1 block potency for imipramine and LY97241 against three disease-associated variants with prominent gain-of-function effects. We chose imipramine because it is approved for pediatric use and has a relatively low proarrhythmia profile in children^43^ even though it can block hERG channels *in vitro*.^44^ We selected LY97241 as the most potent KCNH1 inhibitor reported in the literature. Both compounds were tested at a range of concentrations against WT KCNH1, two recurrent disease-associated variants (R357Q, I494V), and one non-recurrent disease-associated (G375R). We did not select the recurrent variant G496R because whole cell current density in the homozygous state was too low to conduct reliable pharmacological inhibition experiments.

Both imipramine and LY97241 inhibited WT and variant KCNH1 channels, but LY97241 was substantially more potent (for WT KCNH1: imipramine IC_50_ = 1.9 ± 1.1 µM; LY97241 IC_50_ = 0.024 ± 0.005 µM). The recurrent variant I494V exhibited 6-fold more potent block by imipramine than WT channels (I494V: IC_50_ = 0.3 ± 0.1 µM, vs WT: IC_50_ = 1.9 ± 1.1 µM; **Fig. 7**). LY97341 inhibited two variants (R357Q, G375R) with approximately 2-fold greater potency than WT channels (**Fig. 7**). These findings suggest the possibility of variant-specific pharmacology for KCNH1.

**Figure 7.**
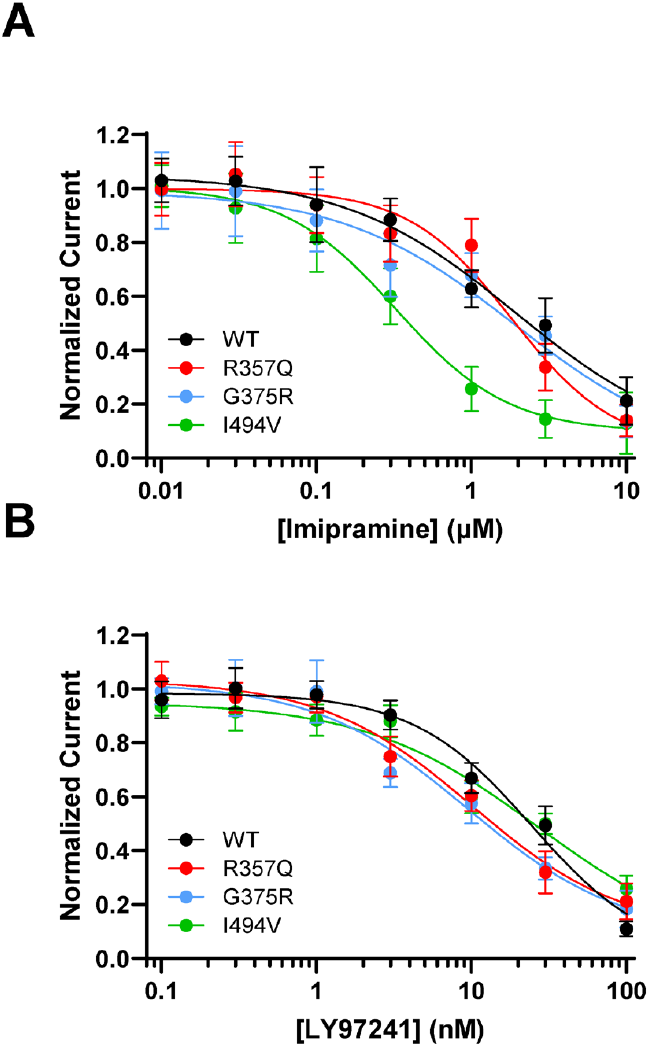
Pharmacology of disease-associated KCNH1 variants. (**A**) Concentration response curves for imipramine inhibition of WT (1.9 ± 1.1 µM), R357Q (1.8 ± 0.6 µM), G375R (1.9 ± 0.9 µM), and I494V (0.3 ± 0.1 µM) in KCNH1 isoform 1 expressed in the homozygous state. (**B**) Concentration response curves for LY97421 inhibition of WT (24.5 ± 5.0 nM), R357Q (10.2 ± 5.3 nM), G375R (9.5 ± 6.2 nM), and I494V (26.7 ± 18.3 nM) in KCNH1 isoform 1 expressed in the homozygous state. All IC_50_ values are reported as mean ± 95% CI (*n* = 5-50).

## Discussion

In this study we determined the functional consequences of several *KCNH1* variants associated with clinically diverse neurodevelopmental disorders. From our data, we draw three main conclusions: 1) most disease-associated *KCNH1* variants exhibit features consistent with gain-of-function including a hyperpolarized activation voltage-dependence, accelerated activation kinetics, or both; 2) KCNH1 gain-of-function drives hyperpolarization of the resting membrane potential in heterologous cells and this is best correlated with the hyperpolarized activation voltage-dependence; 3) KCNH1 alternative splicing can impact the functional properties of disease-associated variants. In addition, we demonstrated that the non-selective potassium channel inhibitors imipramine and LY97241 exhibit higher potency for specific disease-associated KCNH1 variants. Overall, our findings contribute to understanding the molecular pathophysiology of KCNH1-related disorders and support further work to explore the potential therapeutic benefits of drug repurposing for these conditions.

Our approach to investigating KCNH1 variant function is distinct from prior studies of this gene. We exclusively used automated patch clamp, which was used successfully to investigate multiple channelopathies.^27,28,45-51^ Automated patch clamp recording offers several advantages over conventional manual patch clamp recording including the capability to collect much larger sample sizes that boost statistical power, avoidance of operator bias in selecting cells for recording that can skew results in an uncontrolled manner, direct comparisons of WT and variant channels recorded in parallel, and higher throughput allowing study of many more variants.^26^ Additionally, this approach affords the ability to pivot to pharmacological studies with multiple drugs, drug concentrations and multiple variants. Importantly, we studied all KCNH1 variants in the clinically relevant heterozygous state.

Our findings on a subset of disease-associated *KCNH1* variants aligns with data from two prior studies. Kortum *et al*.,^12^ reported functional studies of four individual variants (G375R, L379V, I494V, G496R) and one compound variant (S352Y/V383L) expressed in CHO cells and *Xenopus* oocytes. All variants except G496R exhibited a hyperpolarized activation voltage-dependence and faster activation kinetics (G375R, L379V, I494V only) when expressed in isoform 1 without co-expressed WT KCNH1 (e.g., homozygous state), whereas cells expressing G496R showed no measureable current above background levels unless co-expressed with the WT channel (e.g., heterozygous state). Similarly, Simons *et al*.,^13^ reported hyperpolarized activation for three other variants in addition to I494V expressed in isoform 1 in the homozygous state using HEK293 cells and *Xenopus* oocytes with comparisons to WT channels that was constructed in isoform 2. Collectively, the observations that disease-associated *KCNH1* variants share hyperpolarized activation voltage-dependence appears robust.

However, not all KCNH1 variants we studied exhibited gain-of-function or demonstrable dysfunctional properties. Most notable were K199R and V713E for which we were unable to detect any abnormal functional parameter after testing in both splice isoforms. Neither of these variants appear in gnomAD indicating they are both ultra-rare variants consistent with being pathogenic. However, V713E is classified as likely benign in ClinVar probably based on the reported inheritance from an unaffected parent.^19^ We also observed two variants (R520Q, R520W) that exhibited significant effects on multiple functional parameters in the homozygous state but no differences in the heterozygous state. One of these variants (R520W) is associated with a milder phenotype (normal muscle tone, mild developmental delay, **Table 1**) raising the possibility of lower expressivity. The other variant of this codon (R520Q) was reported in ClinVar without detailed phenotype information. We observed loss-of-function behavior for only one variant (V569M) previously identified in an autism cohort^17^ suggesting a possible function-phenotype divergence.

While the majority of disease-associated variants exhibited biophysical defects consistent with channel gain-of-function, the physiological consequence may influence passive neuronal properties that may dampen neuronal excitability. A key finding from our study that contributes to understanding the pathogenesis of *KCNH1*-related disorders is our discovery that cells expressing heterozygous variants exhibited a hyperpolarized RMP. This observation can be explained by greater potassium ion conductance driving the membrane potential to more negative voltages. We speculate that this could be a major factor causing dysfunction of neural circuits responsible for seizures and possibly other neurological manifestations in *KCNH1*-related disorders. If neurons expressing gain-of-function *KCNH1* variants indeed have a hyperpolarized RMP, then this will likely affect firing threshold. Abnormal membrane potential may also affect cell proliferation and maturation driven by membrane potential in non-excitable cells.^52^ Future experiments using cultured neurons or acute brain slices from genetically engineered animal models may contribute to understanding how KCNH1 gain-of-function disrupts neuronal firing behaviors and circuit excitability. Some features of TBS and ZLS may also be the consequence of KCNH1 dysfunction affecting primary cilia.^53^

Treatment of *KCNH1*-related disorders is not standardized beyond conventional approaches for seizure control and behavioral modulation. Knowing that most *KCNH1* variants exhibit gain-of-function raises the possibility that channel inhibition may have therapeutic benefits. Unfortunately, there are no known KCNH1-specific blockers. However, multiple existing drugs and some investigational compounds are potent inhibitors of this potassium channel and might be therapeutic candidates.^36-41^ A major drawback of targeting KCNH1 is the high likelihood that drugs will block the related hERG channel (encoded by *KCNH2*) expressed in heart. Inadvertent inhibition of hERG is a leading reason for drug withdrawals from the market due to proarrhythmia.^54-56^ It is possible that some KCNH1 variants may exhibit enhanced sensitivity to block by existing drugs that could be effective at lower doses providing a better therapeutic window. In our experiments, we observed that the tricyclic antidepressant drug imipramine exhibits greater potency to block one recurrent KCNH1 variant (I494V) supporting this concept. Neurotoxins that shift the activation voltage-dependence to more depolarized potentials has been proposed as a potential precise therapeutic approach.^11^ The challenge of finding or creating KCNH1-selective compounds should prompt consideration of alternative therapeutic strategies such as allele-specific antisense oligonucleotides or gene editing that avoid off-target hERG inhibition.

### Study limitations

While investigating the functional consequences of ion channel variants in heterologous cells is the standard practice in the field, the physiological impact of dysfunctional KCNH1 variants on native neurons and intact neural circuits will require other cellular or animal model systems that are beyond the scope of our study. Assessing the biophysical behavior of KCNH1 variants in heterologous cells may not be able to predict defects in targeting and localization in native neurons that could be important determinants of pathogenesis even for variants that show no overt functional abnormalities in HEK293 cells (e.g., K199R, V713E). We also did not investigate how KCNH1 variants respond to intracellular signaling molecules that may be important in specific physiological conditions.^57-60^

## Conclusion

This study demonstrated the functional consequences of several previously identified and five unpublished *KCNH1* variants associated with rare neurodevelopmental disorders including TBS and ZLS. Disease-associated variants most often exhibited features consistent with gain-of-function that are distinct from the functional behavior of six rare nonsynonymous population variants having WT channel activity and one variant reported from an autism cohort that exhibited loss-of-function. KCNH1 gain-of-function variants promote hyperpolarization of the RMP, and this may be a driver of abnormal neuronal and neural circuit excitability. We observed allele-specific differences in potency for inhibition by imipramine and an investigational channel blocker (LY97241) for two variants. Our study contributes to understanding the pathogenesis of KCNH1-related disorders and identifies common functional perturbations that may be amenable to pharmacological therapies.

## Acknowledgements

The authors thank the families affected by KCNH1-related disorders for participating in this research study. We are especially grateful to Noreen Haider for compiling a database of families that prompted this study. The authors also thank Dr. Luis Pardo for helpful discussions. We acknowledge the University of Maryland Brain and Tissue Bank through the NIH NeuroBioBank as providing tissue samples.

## Materials and Methods

### Informed consent

Study participants were consented for inclusion in a patient registry established by the Cure KCNH1 Foundation. Informed consent was conducted using a mechanism approved by the Northstar Review Board (Protocol NB00052; learningirb.org). Deidentified clinical and genetic data were accessed by investigators at Northwestern University under an approved data use agreement. Additional study participants were consented using an informed consent mechanism approved by the Institutional Review Board of the Ann and Robert H. Lurie Children’s Hospital of Chicago.

### Cell culture

HEK293T cells (CRL-3216, American Type Culture Collection, Manassas, VA, USA) were maintained in Dulbecco’s modified Eagle’s medium (GIBCO/Invitrogen, San Diego, CA, USA) supplemented with 10% fetal bovine serum (Atlanta Biologicals, Norcross, GA, USA), 2 mM L-glutamine, 50 units/mL penicillin, and 50 µg/mL streptomycin at 37°C in 5% CO_2_.

### Plasmids and mutagenesis

Full-length cDNAs encoding human WT (transcript isoform 1, NCBI RefSeq NM_172362; generously provided by Dr. Roland Schönherr) or variant KCNH1 (non-italicized gene name refers to the cDNA) were engineered in plasmid vectors having a high efficiency encephalomyocarditis virus internal ribosome entry site (IRES) with A6 bifurcation followed by CyOFP (WT vector) or EGFP (variant vectors) as described previously.^25^ A single silent nucleotide change (c.894T>C) was made in the KCNH1 coding region to disrupt a potential prokaryotic promoter sequence to stabilize the plasmid during growth in bacteria. Some variants and WT were also made in KCNH1 transcript isoform 2 (NCBI RefSeq NM_002238), which we constructed by site-directed mutagenesis.

Variants were introduced into KCNH1 by site-directed mutagenesis using Q5 2X high-fidelity DNA polymerase Master Mix (New England Biolabs, Ipswich, MA, USA) as previously described.^25^ Mutagenic primers (**Supplementary Table S1**) were designed using custom software (Q5 Designer; available at https://doi.org/10.18131/9tpj3-bx112). Full-length plasmids were sequenced using nanopore sequencing technology (Primordium Laboratories, Arcadia, CA, USA or Plasmidsaurus, Lexington, KY, USA) and analyzed using a custom multiple sequence alignment tool (Multiple Sequence Iterative Comparator [MuSIC], available at https://doi.org/10.18131/h6hc6-n0j20). Only plasmids with perfect matches to the expected sequence were used for experiments. Endotoxin-free plasmid DNA was purified using Nucleobond Xtra Maxi EF columns (Macherey-Nagel Inc., Allentown, PA, USA) and re-suspended in endotoxin free water.

### Electroporation

For automated electrophysiology experiments, plasmids encoding WT and/or variant *KCNH1* were electroporated into HEK293T cells using the MaxCyte STX system (MaxCyte Inc., Gaithersburg, MD, USA).^26^ Cells were grown to 70-80% confluence and harvested using TrypLE (ThermoFisher, Waltham, MA, USA). A 500 μL aliquot of cell suspension was used to determine cell number and viability using an automated cell counter (ViCell, Beckman Coulter, Brea, CA, USA). Remaining cells were collected by gentle centrifugation (193 x g, 4 min) at room temperature, followed by washing the cell pellet with 5 mL electroporation buffer (EBR100, MaxCyte Inc.). Cells were resuspended at a final density of 10^8^ viable cells/mL.

To examine functional properties in the homozygous state, 45 µg of WT or variant KCNH1 plasmid DNA was mixed with 100 µL of cell suspension (10^8^ cells/mL) for electroporation. To examine functional properties of co-expressed WT and variant channels (heterozygous state), 40 µg total plasmid DNA (20 µg WT and 20 µg variant) was electroporated. For all electroporations, the DNA-cell suspension mix was transferred to an OC-100x2 processing assembly (MaxCyte, Inc.) and electroporated using the Optimization 4 protocol. Immediately after electroporation, 10 µL recombinant human deoxyribonuclease I (dornase alpha, 1 mg/mL, Genentech, San Francisco, CA, USA) was added to the DNA-cell suspension mix, and the entire mixture was transferred to a 60 mm dish and incubated for 30 minutes at 37°C in 5% CO_2_. Following incubation, cells were gently resuspended and grown in a T75 flask for 48 hours at 37°C in 5% CO_2_. Cells were then harvested, transfection efficiency determined by flow cytometry (see below), and then frozen in 1 mL aliquots at 1.8 x 10^6^ viable cells/mL.

Transfection efficiency following electroporation was assessed prior to cell freezing using a benchtop flow cytometer (CytoFLEX, Beckman Coulter). Forward scatter (FSC), side scatter (SSC), green (EGFP) and red (CyOFP) fluorescence were recorded. FSC and SSC were used to gate single viable cells and eliminate doublets, dead cells, and debris. Ten thousand events were recorded for each sample, and non-transfected HEK293T cells were used as a negative control to set gates. A 488 nm excitation laser was used for green fluorescence for experiments examining channels in the homozygous state. For co-expression experiments, a 488 nm laser was used to excite both green and red fluorescence. A compensation matrix was generated using HEK293T cells expressing only single fluorescent markers and applied to co-transfected cells to account for spectral overlap. The percentage of co-transfected cells was determined from plots of green vs red fluorescence intensity.

### Cell preparation for automated electrophysiology

Electroporated cells were thawed one day before experiments and grown ∼24 hours at 37^°^C in 5% CO_2_. Prior to experiments, cells were dispersed using TrypLE, and a 500 µL aliquot was taken to determine cell number and viability by automated cell counting. Cells were gently centrifuged (193 × g) for 4 min at room temperature and resuspended at a final density of 180,000 viable cells/mL in external recording solution (see below). Cells were allowed to recover on a shaking rotating platform (200 rpm) at 15°C for 30-45 minutes prior to recording.

### Automated electrophysiology

Automated voltage-clamp and current clamp recordings were performed using the Nanion SyncroPatch 384 platform (Nanion Technologies, Munich, Germany) using medium resistance chips as described previously.^27,28^ The composition of the external solution contained (in mM): 140 NaCl, 4 KCl, 2 CaCl_2_, 1 MgCl_2_, 10 HEPES, 5 glucose, with the final pH adjusted to 7.4 with NaOH, and osmolality adjusted to 300 mOsm/kg with sucrose. The composition of the internal solution was (in mM): 60 KF, 50 KCl, 10 NaCl, 20 EGTA, 10 HEPES, with the final pH adjusted to 7.2 with KOH, and osmolality adjusted to 300 mOsm/kg with sucrose. High resistance seals were obtained by addition of a 3:1 mixture of external solution and seal enhancer solution comprised of (in mM): 125 NaCl, 3.75 KCl, 10.25 CaCl_2_, 3.25 MgCl_2_, 10 HEPES, final pH adjusted to 7.4 with NaOH. Prior to recording, cells were washed twice with external solution, and the final concentrations of CaCl_2_ and MgCl_2_ were 3 mM and 1.3 mM, respectively. Unless otherwise noted, chemicals were obtained from SigmaAldrich (St. Louis, MO, USA).

For whole-cell voltage clamp recording, cells were held at -80 mV, followed by a 1 sec pulse from -110 to +50 mV in 10 mV increments. Tail currents were measured during a 300 ms pulse to -30 mV. Resting membrane potential was measured in by current clamp using ten 1 sec sweeps with zero direct current injection immediately after obtaining whole-cell configuration. Only cells with seal resistance >500 MΩ, series resistance <20 MΩ, and cells capacitance >2 pF were used for final data analyses. For additional quality control, only cells with current amplitude >250 pA at +50 mV, a minimum baseline current ≥ -200 pA, and a minimum tail current from the +50mV test pulse ≥ 50 pA were used for final analysis.

### Electrophysiological data analysis

Data were analyzed and plotted using a combination of DataControl384 (Nanion Technologies), Clampfit 10.4 (Molecular Devices, San Jose, CA, USA), Microsoft Excel (Microsoft, Redmond, WA, USA), and GraphPad Prism (GraphPad Software, San Diego, CA, USA). Whole-cell currents were normalized to membrane capacitance to calculate current density. To assess voltage-dependence of activation, currents at steady state were converted to conductance and GraphPad Prism was used to fit voltage-dependence of activation data to a Boltzmann function to determine V½ (voltage at which 50% of the channels are open) and *k* (slope factor). Activation time constants were determined using Clampfit10.4 from a single exponential fit of the current recorded at +10 mV recorded at the beginning of the test pulse through to the steady-state phase. Data were normalized to WT run in parallel and data were treated as not normally distributed for statistical analyses. Unless otherwise noted, data are presented as mean ± 95% confidence interval (CI). Statistical analyses to determine differences between WT and variant KCNH1 were performed using a Kruskal-Wallis test followed by Dunn’s post-hoc test for multiple comparisons. The threshold for statistical significance was *P* < 0.02. Exact *P*-values are presented in the supplementary dataset files.

### Pharmacology

Automated patch clamp was used to assess pharmacological properties of WT, R357Q, G375R (G>A), and I494V. Imipramine and LY97124 were prepared as 10 mM stock solutions in DMSO and serially diluted in external solution for concentration response experiments. Final drug containing solutions contained 0.1% DMSO. Concentration response experiments were performed in parallel with vehicle (0.1% DMSO) treated cells. For concentration response experiments, currents were elicited by a 2 sec test pulse to +50 mV from a holding potential of -80 mV. Baseline currents were recorded for 1 min, followed by a 5 min application of vehicle, imipramine, or LY97241. Inhibition was calculated by comparing the current level remaining after drug exposure to the current level remaining in vehicle treated cells. Concentration response curves were fit with a four-parameter Hill equation using GraphPad Prism. Data are reported as mean ± 95% CI, and IC_50_ values are reported as IC_50_ ± 95% CI. Imipramine was obtained from SigmaAldrich and LY97241 was synthesized by the Northwestern University Center for Molecular Innovation and Drug Discovery.

### *KCNH1* splice isoform expression in human brain

Post-mortem human brain tissues were obtained from University of Maryland Brain and Tissue Bank through the NIH NeuroBioBank. Frozen primary motor cortex (Broadman area 4) from donors ranging in age from 18 post conception weeks (pcw) to 20 years were used. Total RNA was extracted with TriZol reagent (ThermoFisher Scientific, Waltham, MA, USA) and cDNAs were synthesized using SuperScript IV First-Strand synthesis system with ezDNase (ThermoFisher) according to manufacture protocols. Regions spanning alternative splicing sites were amplified by PCR using Q5 DNA polymerase (primer pair provided in **Supplementary Table S1**). Purified amplicons were analyzed by next-generation sequencing (GENEWIZ from Azenta Life Sciences, South Plainfield, NJ, USA). Sequencing reads were aligned to the reference sequences of predicted splicing isoforms using CRISPResso2.^29^ Expression of each splice isoform was quantified as a percentage of total aligned amplicons and presented as mean ± SEM. The contribution of each isoform within each age group was compared by unpaired t-test. The ratio of isoform 1/isoform 2 among age groups was compared using a one-way ANOVA followed by Tukey’s post-test.

## SUPPLEMENTARY MATERIAL

### Supplementary Tables

**Table S1**. Primer sequences

### Supplementary Figures

**Fig. S1**. Average whole-cell current traces of *KCNH1* variants in the homozygous state

**Fig. S2**. Average whole-cell current traces of *KCNH1* variants in the heterozygous state

**Fig. S3**. Expression of *KCNH1* splice isoforms in human brain tissues during development

**Fig. S4**. Comparison of functional properties of *KCNH1* splice isoforms

**Fig. S5**. Average whole-cell current traces of *KCNH1* variants in isoform 2

**Fig. S6**. Correlation between ΔRMP with fold change in activation time constant

**Supplementary Table S1.**
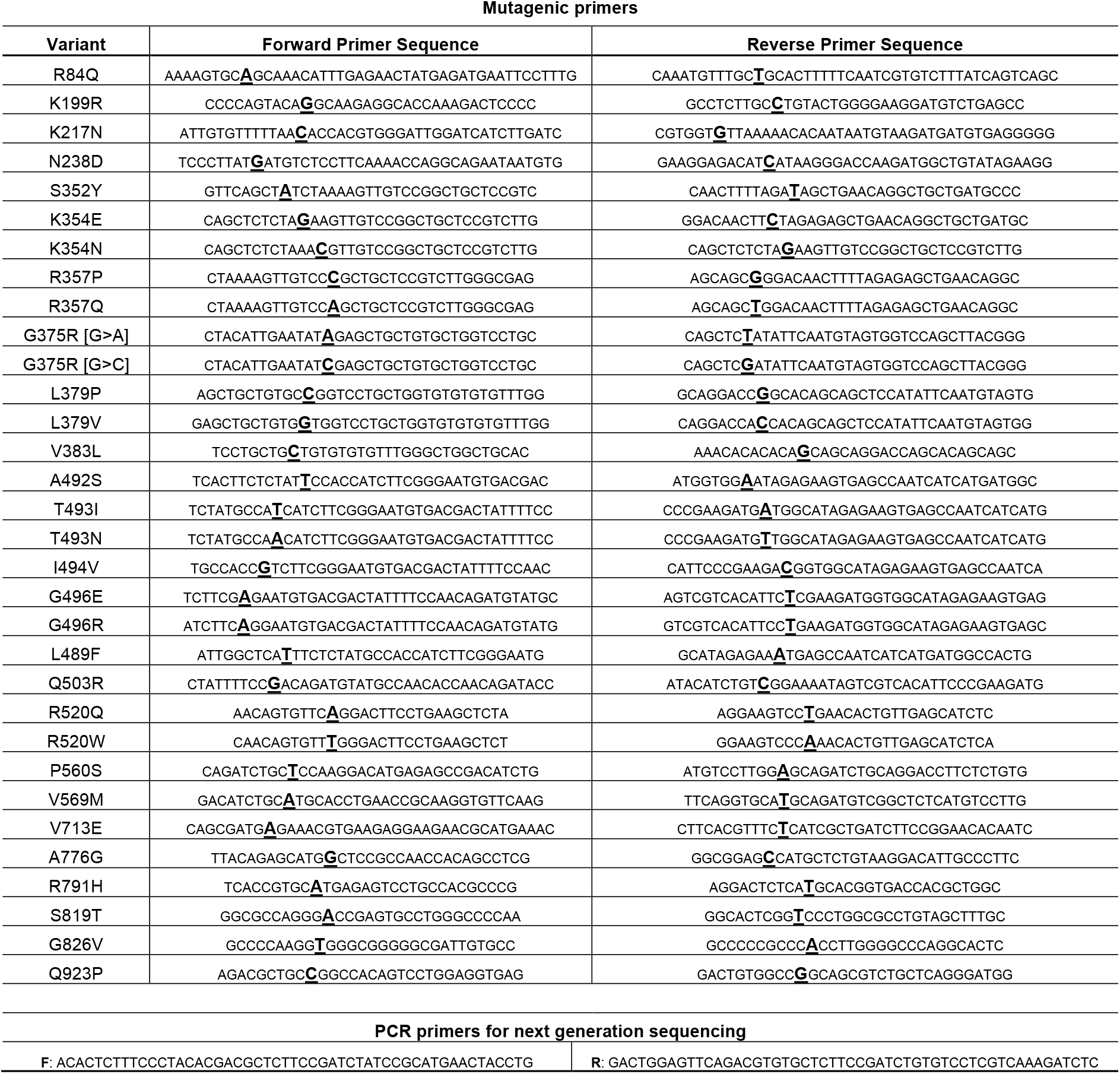
Primer sequences.

**Fig. S1.**
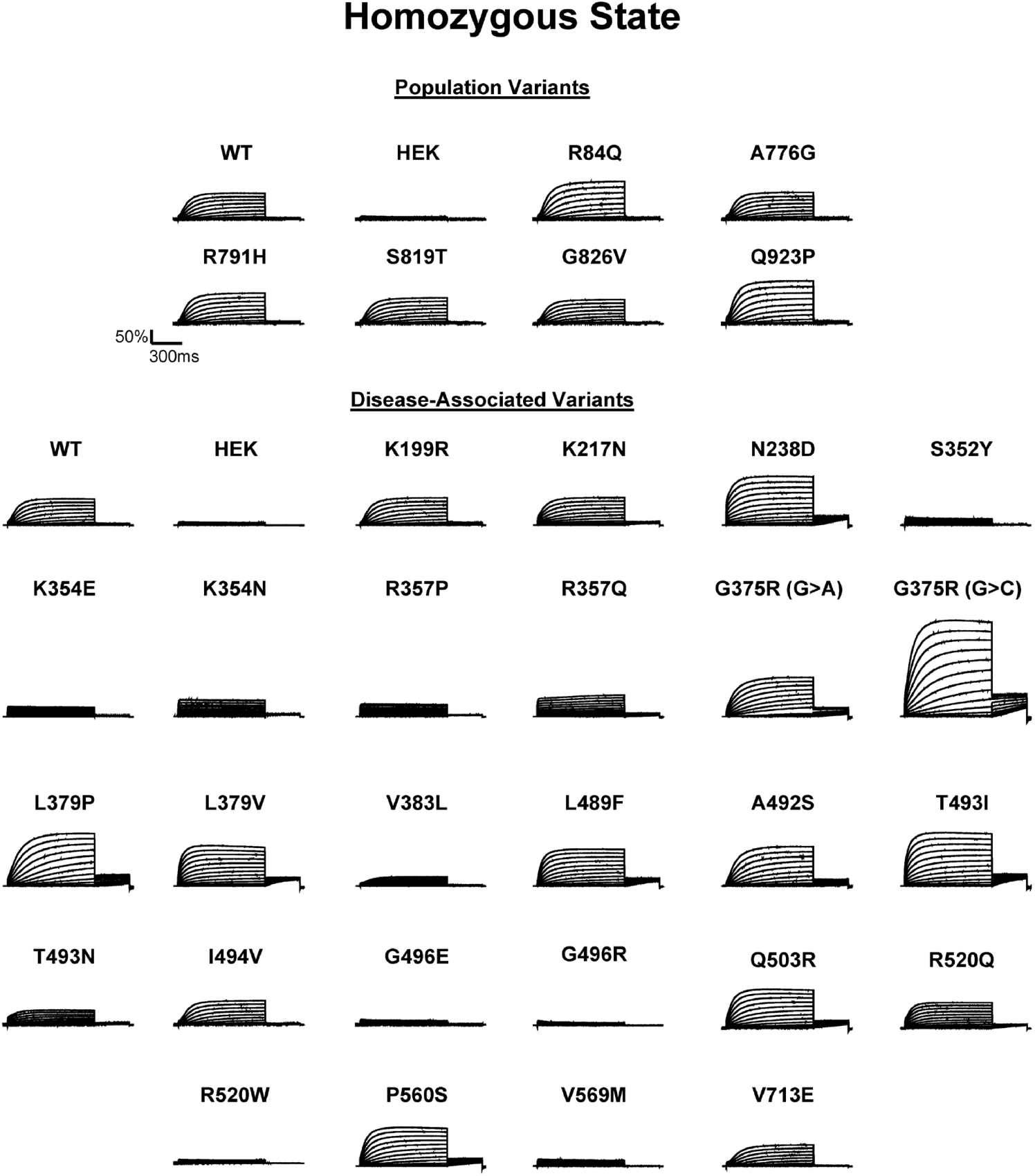
Average whole-cell current traces of KCNH1 variants in the homozygous state. Average normalized whole-cell currents of KCNH1 variants (*n* = 7-57 cells).

**Fig. S2.**
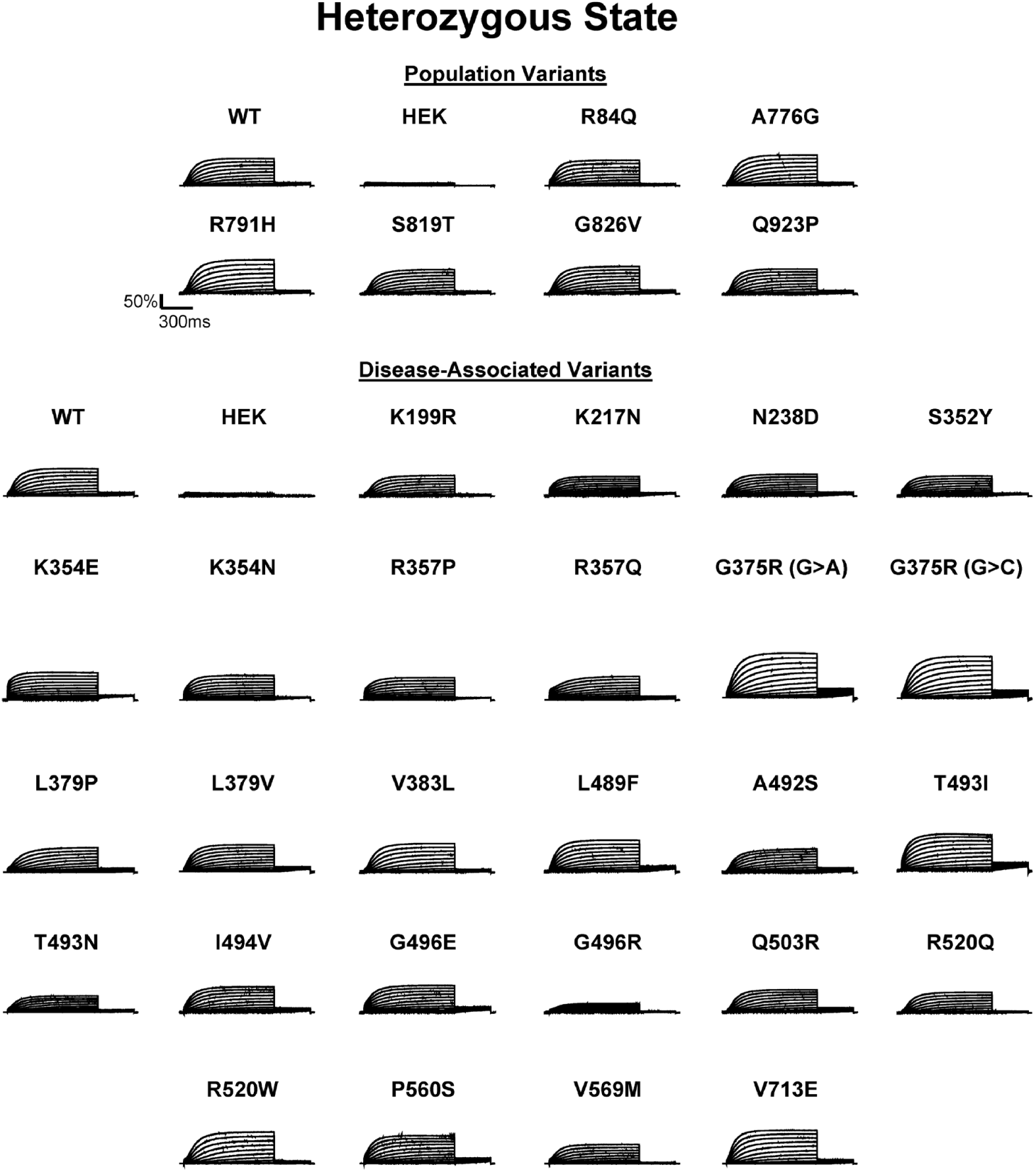
Average whole-cell current traces of KCNH1 variants in the heterozygous state (*n* = 15-94 cells).

**Fig. S3.**
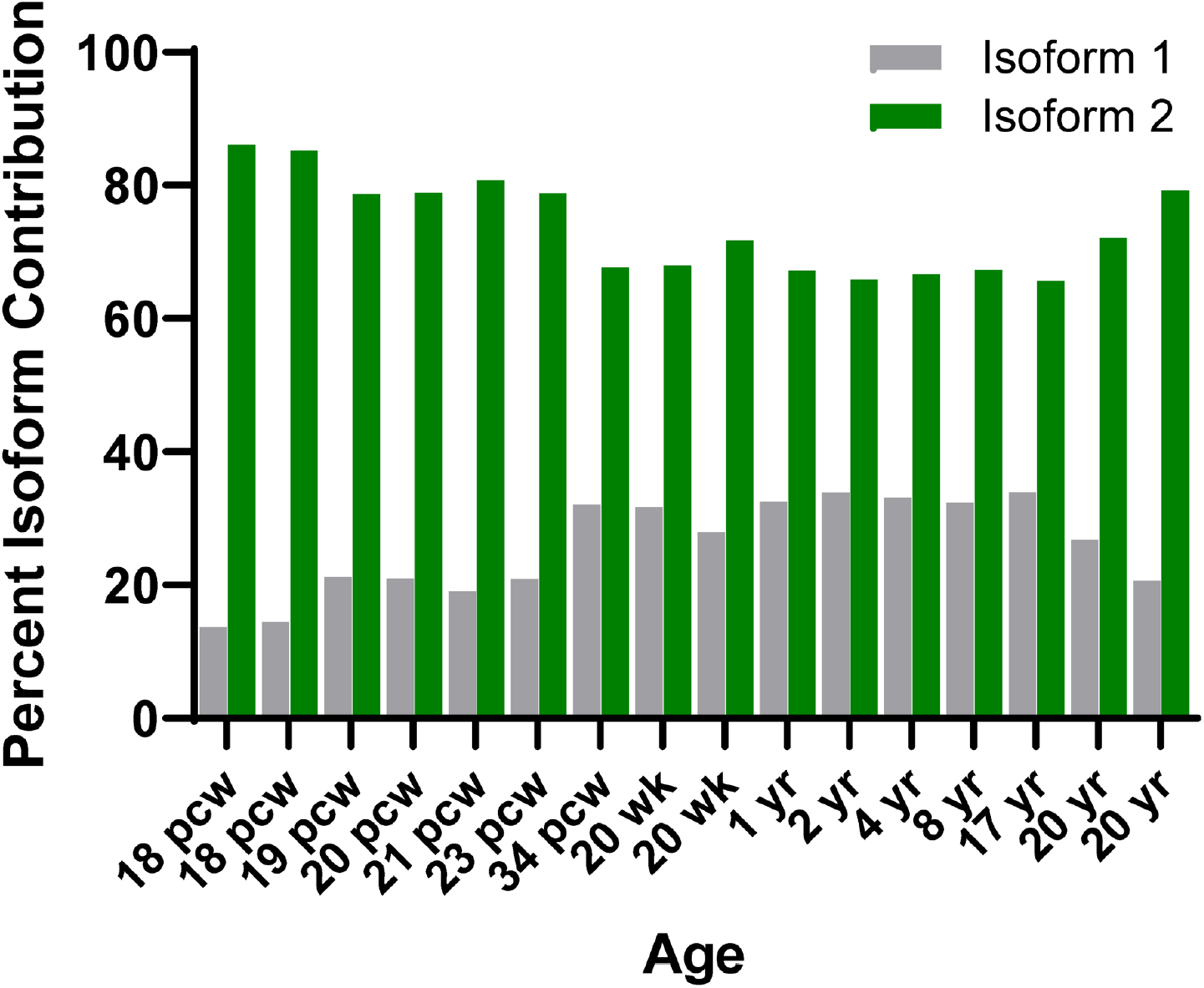
Expression of *KCNH1* splice isoforms in human brain tissues during development. Contribution of each *KCNH1* splice isoform to total *KCNH1* message from individual human brain tissue samples ranging18 post-conception week (pcw) to 20 years of age (*n* = 2-7 samples).

**Fig. S4.**
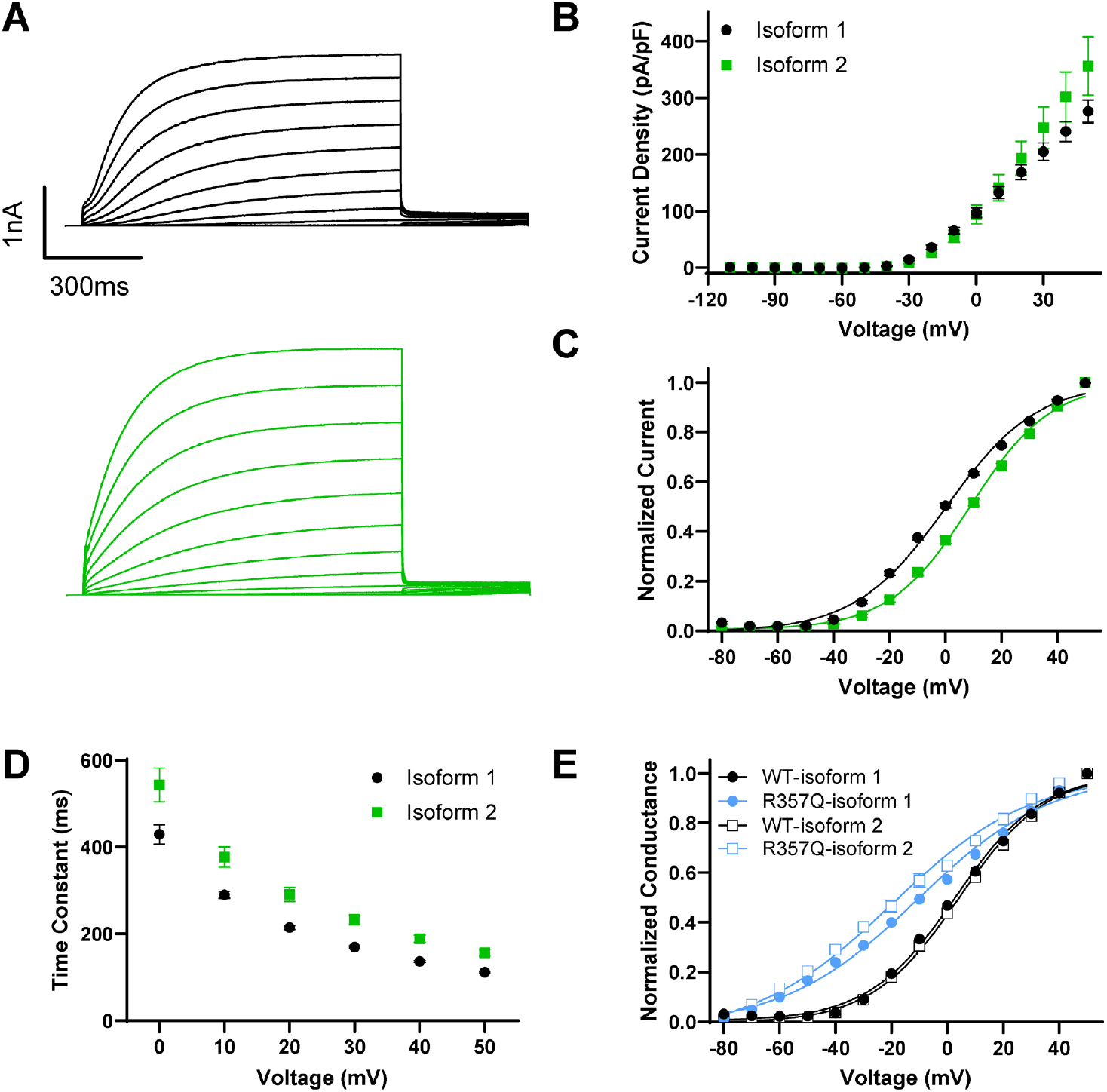
Comparison of functional properties of KCNH1 splice isoforms. **(A)** Average whole-cell current traces for WT isoform 1 (black) and isoform 2 (green). Summary data showing the **(B)** average current-voltage relationship, **(C)** conductance-voltage relationship, and **(D)** time constant of activation at 10 mV. **(E)** Average conductance-voltage relationship of WT (black) and R357Q (blue) in isoform 1 (circles) and isoform 2 (squares). All data are plotted as mean ± 95% CI (*n* = 161-504 cells).

**Fig. S5.**
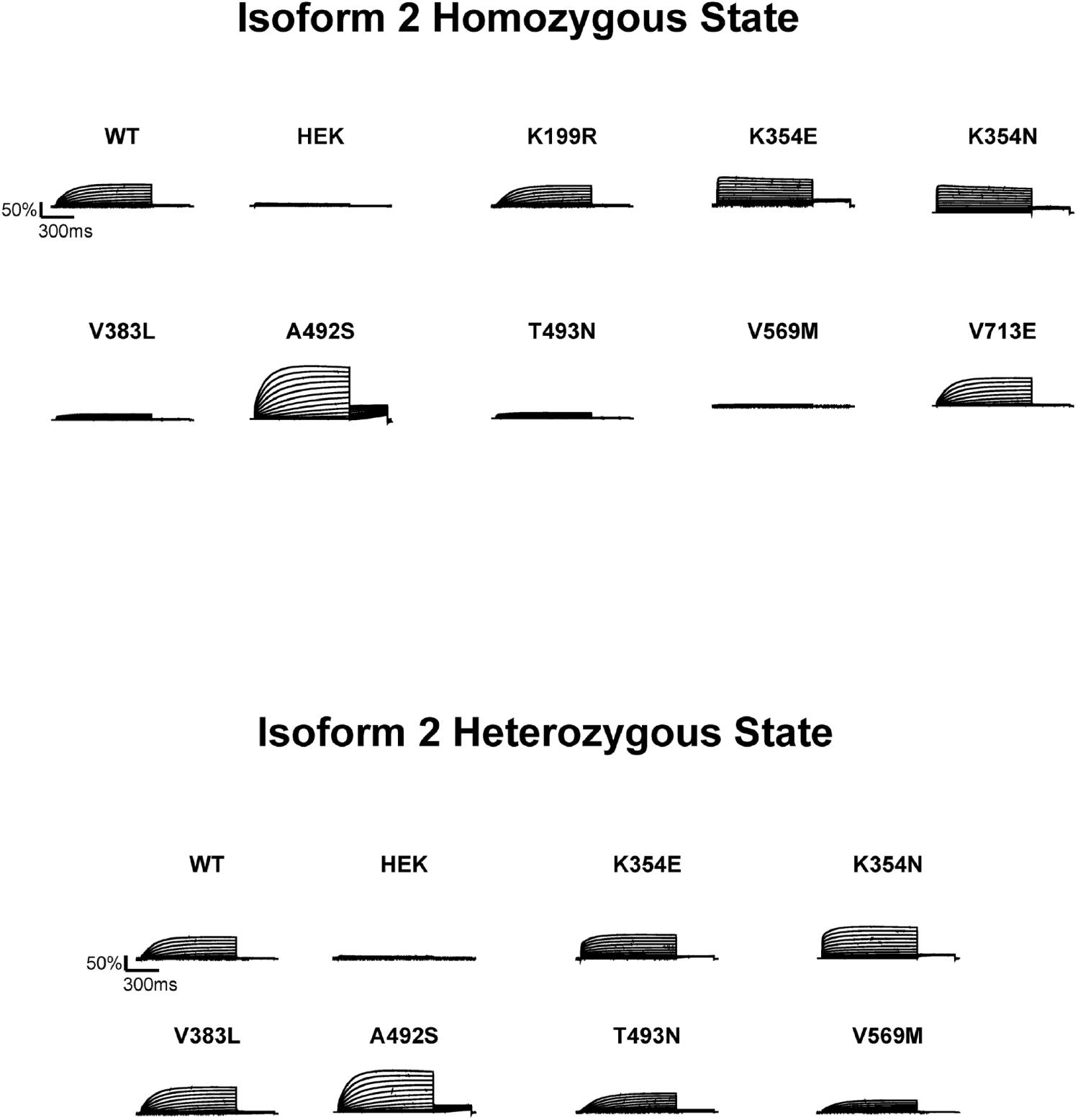
Average whole-cell current traces of KCNH1 variants in isoform 2 (*n* = 14-107 cells).

**Fig. S6.**
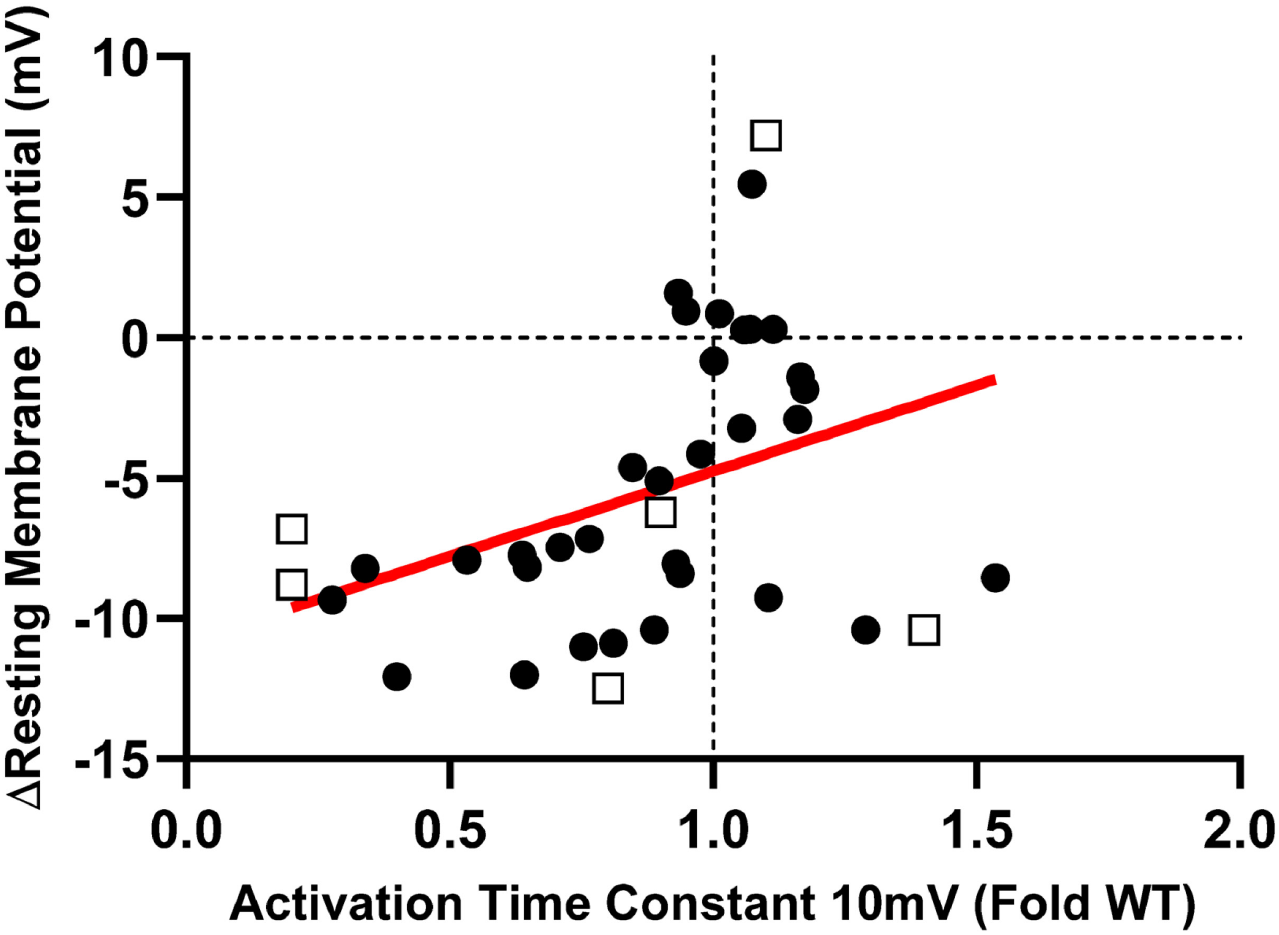
Correlation between ΔRMP with fold change in activation time constant. Correlation between ΔRMP and fold change in activation time constant. Red line is a fitted regression line (*P*=0.0192). Circles represent data obtained from isoform 1, and squares represent data obtained from isoform 2.

